# Reconstruction of phospholipid synthesis by combing in vitro fatty acid synthesis and cell-free gene expression

**DOI:** 10.1101/2021.08.03.454925

**Authors:** Sumie Eto, Rumie Matsumura, Mai Fujimi, Yasuhiro Shimane, Samuel Berhanu, Takeshi Kasama, Yutetsu Kuruma

**Affiliations:** Earth-Life Science Institute, Tokyo Institute of Technology, 2-12-1 Ookayama, Meguro-ku, Tokyo 152-8550, Japan; Institute for Extra-cutting-edge Science and Technology Avant-garde Research (X-star), Japan Agency for Marine-Earth Science and Technology (JAMSTEC), 2-15 Natsushima-cho, Yokosuka 237-0061, Japan; PRESTO, Japan Science and Technology Agency (JST), 4-1-8 Honcho Kawaguchi, Saitama 332-0012, Japan

**Keywords:** Artificial cell, self-reproduction, lipid synthesis, Cell-free system, membrane division

## Abstract

Phospholipid synthesis is a fundamental process that promotes cell propagation and, presently, is the most challenging issue in artificial cell research aimed at reconstituting living cells from biomolecules. Here, we constructed a cell-free phospholipid synthesis system that combines *in vitro* fatty acid synthesis and a cell-free gene expression system that synthesizes acyltransferases for phospholipid synthesis. Fatty acids were synthesized from acetyl-CoA and malonyl-CoA, then continuously converted into phosphatidic acids by the cell-free synthesized acyltransferases. Because the system can avoid the accumulation of synthetic intermediates that suppress the reaction, the yield of phospholipid has significantly improved from previous schemes (up to 400 µM). Additionally, by adding enzymes for recycling CoA, we synthesized phosphatidic acids from acetic acid and bicarbonate as carbon sources. The constructed system is available to express the genes from pathogenic bacteria and to analyze the synthesized phospholipids. By encapsulating our system inside giant vesicles, it would be possible to construct the artificial cells in which the membrane grows and divides sustainably.

## Introduction

Recent attempts to reconstitute cellular functions aim, ultimately, artificial construction of a whole living cell ^1^ to understand the basic principles of life phenomenon. Some cell machinery such as gene expression ^2^, genome replication ^3, 4^, cell division ^5^, and energy generation ^6^ has been reconstructed partially or totally by assembling the responsible biomolecules. Most of them can also function inside lipid membrane vesicles artificially prepared in cell size, and such a system is called an artificial cell. However, a reproduction of cell membrane growth and division, which is necessary for achieving cell proliferation, remains a challenge.

So far, several reports have demonstrated membrane growth and division using noncanonical lipids. Kurihara *et al*. converted the amphiphile precursor, which taken from the outside of the vesicles, into the constituent lipid employing a catalyst that pre-existed in the vesicle membrane ^7, 8^. And Devaraj’s group synthesized *de novo* phospholipids on the vesicle membrane by conducting the bind between acyl-CoA (or acyl-AMP) synthesized by FadD enzyme and lysolipids activated through the insertion of amine^9, 10^. Although these chemically produced lipids do not exist *in vivo*, it proved that membrane growth and division can be reproduced as a physicochemical reaction when phospholipids were newly synthesized on the lipid bilayer. Interestingly, the same principle can be found in L-form type bacteria, which lack a cell wall but can propagate in an FtsZ-independent manner through overproducing phospholipids ^11^.

In more biological approaches, some attempts have reported the enzymatic synthesis of canonical phospholipids. We have previously demonstrated to synthesize phospholipids by synthesizing acyltransferases inside liposomes encapsulating those responsible genes and a cell-free protein synthesis system^12^. Using the same approach, Danelon’s group have synthesized various phospholipids found in living cell membranes by cell-free synthesized phospholipid synthases ^13, 14^. However, in both studies, the amount of synthesized phospholipid was not sufficient for the deformation of the vesicle membrane. Synthesis of fatty acid, which bearing the volume of the lipid bilayer of vesicle membrane, has also been reconstructed by Yu *et al*. ^15^ but the yields arrested again after reaching the plateau at 200-300 µM (100-150 µM as diacyl-phospholipid). While these results clearly show that we can newly synthesize phospholipids if the enzymes are available on the membrane, they also defined that the difficulty of increasing the yield of phospholipid synthesis is the core of the problems in artificial cell research. For making double the vesicle surface area with a diameter of 30 µm, for example, 1mM phospholipids must be synthesized inside as new lipids (Supplementary Table1). And even more difficult is that those all must insert into the membrane. Although it is possible to feed fatty acids from the outside of the vesicles, they need to eventually convert to more stable phospholipids because of the poor physical stability of the fatty acid membrane. To overcome this problem, it is crucial to build an *in vitro* system capable of synthesizing phospholipids more efficiently and sustainably. In this study, we designed and constituted a new cell-free system capable of producing phospholipids in sub-millimolar by combining a fatty acid synthesis system and a phospholipid synthesis system (Fig. 1A). The phospholipid synthesis system was established by expressing genes of acyltransferases in a cell-free protein synthesis system (PURE system) ^2^. We also implemented the acetyl-CoA and malonyl-CoA syntheses pathway, which uses acetic acid and bicarbonate as carbon sources, by adding three more responsible enzymes. Additionally, we demonstrated that our cell-free system is available to express the genes of acyltransferases from pathogenic bacteria and to evaluate their functions rapidly. We believe that, by installing this cell-free phospholipid synthesis system inside vesicles, the construction of sustainably self-growing and self-dividing artificial cells will be closer to reality.

**Figure 1.**
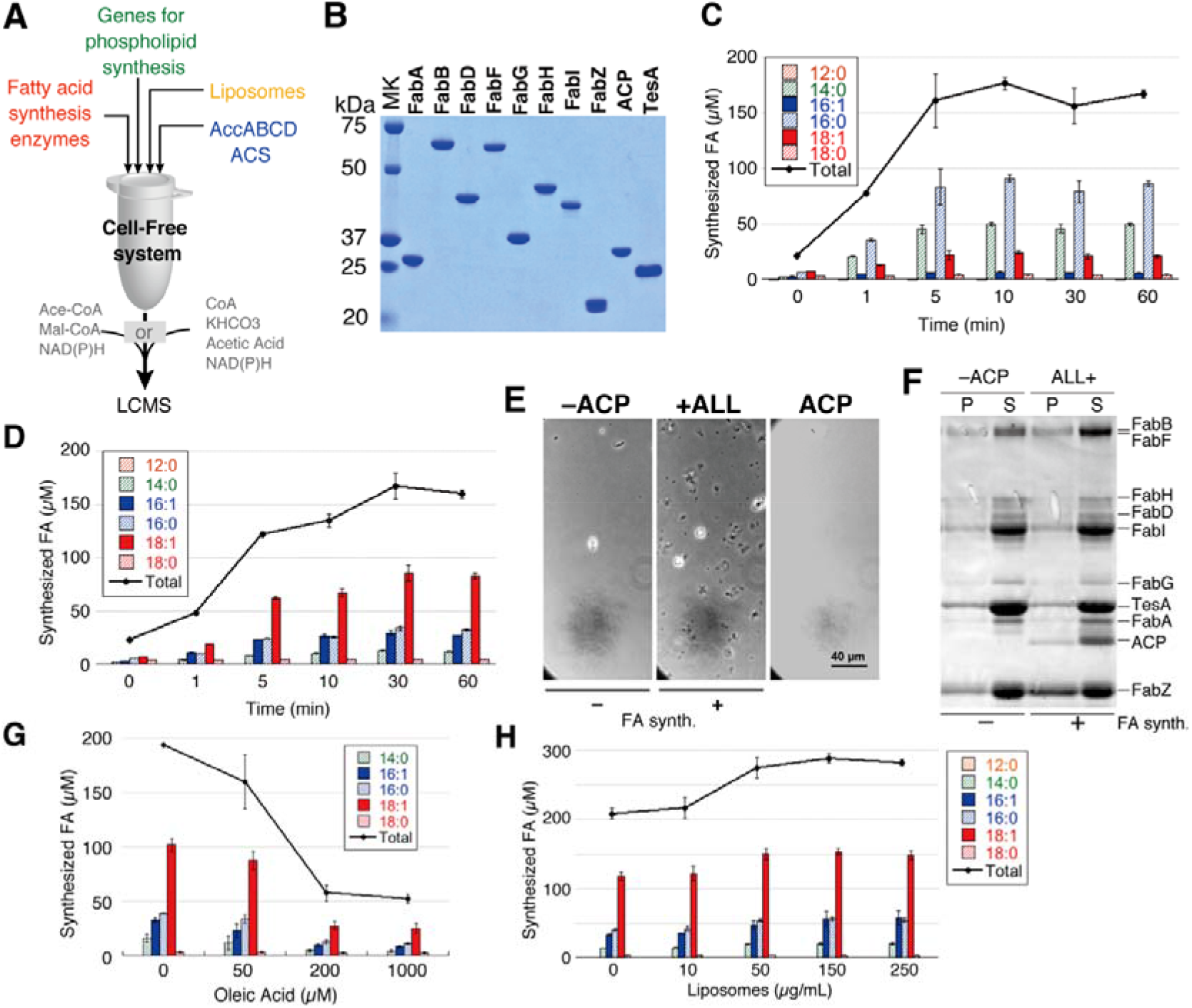
Fatty acid synthesis in a reconstructed cell-free system. (A) Schematic overview of the cell-free phospholipid synthesis system. After the gene expression of phospholipid syntheses in the presence of the enzymes for fatty acid synthesis, liposomes, AccABCD and ACS, phospholipid synthesis reaction is initiated by the addition of substrates for fatty acid (a set of acetyl-CoA and malonyl-CoA, or a set of CoA, KHCO3, and acetic acid) and NAD(P)H. The synthesized fatty acids were quantified by an LCMS. (B) SDS-PAGE image of purified Fab enzymes, ACP, and TesA. The positions of the molecular size are described on the left side of the gel image. (C) Fatty acid synthesis in the reconstituted *in vitro* fatty acid synthesis system containing 10µM FabZ, 1µM FabA, and 1µM FabB. The types of synthesized fatty acids and the total yield are shown in the inset. (D) Fatty acid synthesis is the same as (C), but 1µM FabZ, 10µM FabA and 1µM FabB. (E) Optical microscopy images of the reaction mixture after fatty acid synthesis. +All and −ACP indicate the mixture containing all enzymes and missing ACP, respectively. Error bar: 40 µm. (F) SDS-PAGE analysis of the reacted reaction mixture in the presence or absence of ACP. Each position of the enzymes is described on the right side of the gel image. P and S represent precipitate and supernatant, respectively. (G Inhibition of fatty acid synthesis by fatty acid (oleic acid). (H) Enhancement of fatty acid synthesis by addition of liposomes. Error bars indicate the standard deviation of triplicate measurements. Abbreviations: ACS: acetyl-CoA synthetase, ACP: acyl carrier protein, MK: molecular marker.

## Results

### In vitro fatty acid synthesis

To reconstruct the *in vitro* fatty acid synthesis, we purified eight kinds of fatty acid-binding enzymes (FabA, FabB, FabD, FabF, FabG, FabH, FabI, and FabZ), acyl carrier protein (ACP), and thioesterase 1 (TesA) (Fig. 1B). We confirmed that the purified ACP was modified with 4’-pantetheine to achieve a holo-form (Supplementary Fig. 1). These enzymes were mixed with two substrates (acetyl-CoA and malonyl-CoA) and electron donors (NADH and NADPH) based on a previous report^15^ to start fatty acids synthesis reaction. The synthesized fatty acids were analyzed by an LCMS to quantify each type of fatty acid. Most of the synthesized products were saturated fatty acids, mainly C14:0 and C16:0 (Fig. 1C), and the concentration of total fatty acids reached 150-200 µM in five minutes after starting the reaction. It was consistent with the previous report ^15^.

Next, we tried to change the ratio of saturated and unsaturated fatty acids in the products by tuning the components in the reaction. According to Feng and Cronan ^16^, the production ratio of saturated and unsaturated fatty acids is determined by the balance between FabA and FabZ, sharing the same dehydration step (Supplementary Fig. 2). FabA has a function not only as a dehydratase but also as an isomerase. Therefore, the product synthesized in the FabA rich condition contains an angled *cis* unsaturation in fatty acids. We changed the ratio of FabA:FabZ from 1:10 to 10:1 (mol.). With this, FabB, which specially catalyzes the next round of carbohydrate elongation on an unsaturated fatty acid, was also increased. The result showed that about 70 % of the products were turned into unsaturated fatty acids (Fig. 1D), whereas those were only 15 % when FabA concentration was low. We demonstrated that the constructed system is highly controllable in the production of saturated or unsaturated fatty acids. However, the yield quickly reached a plateau about ten minutes after the reaction and did not increase after that. To find the limiting factors, we tried to observe a recovery of fatty acid synthesis by additionally supplying substrates or electron donors at the point of plateau. Nevertheless, there was no significant increase was found in either case (Supplementary Fig. 3). We also examined fatty acid synthesis in an anaerobic condition, but again no difference was observed (Supplementary Fig. 4).

Interestingly, we found that the reaction mixture became slightly turbid during the reaction. This was because of the appearance of tiny aggregates that could be observed by optical microscopy (Fig. 1E). To know what is aggregating, we precipitated them by centrifugation and analyzed by SDS-PAGE. The results showed that a significant band of FabZ appeared in the precipitate fractions (Fig. 1F), suggesting the most of the aggregates consist of FabZ. The aggregates of FabZ did not appear when the essential enzyme for fatty acid synthesis (e.g. ACP) was absent in the reaction (Fig. 1E and F, Supplementary Fig. 5). These results imply the synthesized fatty acids induced the formation of abnormal protein structure, mostly FabZ.

Given the results of the reaction arrest and the enzyme aggregations, we hypothesized that synthesized fatty acids are inhibiting the production of fatty acids. To verify this hypothesis, we tried to synthesize fatty acid in the presence of fatty acid (18:1, oleic acid) in various concentrations. The result clearly showed that the yield drastically decreased as increasing the concentration of fatty acid; the productivity dropped to 25 % when more than 200µM oleic acid was presented (Fig. 1G). This result indicates that the synthesized fatty acids not only form stable micelles in the solution but also attach to the enzymes and inhibit their activities. Such an unfavourable effect of fatty acids onto proteins is probably due to their molecular nature, similar to surfactants. In fact, fatty acid-induced enzyme inhibition has been known in other protein^17^.

If so, the inhibitory effect might be reduced by supplying stable lipid membranes (e.g. liposomes) that work as a reservoir of the synthesized fatty acids. To test this, we prepared liposomes and introduced them into the reaction mixture during fatty acid synthesis. The result showed that, as we expected, the yield of fatty acid increased; a maximum of 1.5-fold when 150 µg/mL liposomes were added (Fig. 1H). Supplying liposomes tended to suppress the formation of aggregates (Supplementary Fig. 6). And, this suppression was observed even after the appearance of the aggregates, which dissolved by liposomes (Supplementary Fig. 6). These results indicate that the actual concentration of free or micelle fatty acids decreased by being trapped in the liposome membrane, resulting in extended enzyme life and synthesized more fatty acids.

Quantitative analysis shows that when the same amount of fatty acids as the reaction in the bulk was synthesized in a giant unilamellar vesicle (GUV) with a diameter of 30 µm, it corresponds to only 15% of the lipid content of the vesicle. (Supplementary Text1). It is still far from constructing a self-growing-and-dividing GUV. And, even worse, not all the fatty acids localize onto the vesicle membrane, since fatty acids are in equilibrium between free diffusion and aggregate due to their high critical aggregation concentration. We concluded that it is impossible to apply the fatty acid synthesis system to the construction of artificial cells that enable self-reproduction. Thus, we revised the design of the system to increase its productivity.

### Expression of PlsX and PlsY by the cell-free system in balk or in vesicles

In the first version of the system, the synthesized fatty acids are released to the aqueous reaction solution by the function of TesA, therefore inactivated Fab enzymes. But TesA does not exist in the cytoplasm because it is originally periplasmic enzyme ^18^. In bacteria, newly synthesized fatty acids are directly localized onto the cell membrane in the form of fatty acid-ACP (acyl-ACP). On the membrane surface, acyl-ACP is converted into acyl-PO_4_ by PlsX ^19^, sequentially, acyl-PO_4_ binds to glycerol-3-phosphate by the catalytic function of PlsY to form lysophosphatidic acid (LPA) (Fig. 2A) ^19^. If there are both PlsX and PlsY in the system, it can be expected continuous LPA synthesis from fatty acid synthesis, avoiding the risk of enzyme inactivation and poor membrane localization of the synthesized fatty acids. This is also a favourable system when we implement inside GUV, where internally synthesized fatty acids directly integrate into the vesicle membrane through converting to LPA. However, introducing these enzymes is very difficult because these are membrane proteins requiring detergent in the step of purification. Detergents melt vesicles. Thus, the only way to solve this problem is to synthesize both enzymes inside GUV. For syntheses of PlsX and PlsY, we used a reconstructed cell-free system, PURE system ^2^. Both proteins were successfully co-synthesized in the PURE system, in a bulk (test tube) condition (Fig. 2B) in which about 1µM proteins are synthesized (Supplementary Fig. 7). Note that DnaKJE chaperon was supplied in the PURE system because the solubility of PlsX is highly dependent on DnaKJE (http://www.tanpaku.org/tp-esol/index.php?lang=en) ^20^. In the presence of liposomes, more than 70% of the synthesized PlsX and PlsY were collected in the supernatant fractions after centrifugation of the reaction mixture, suggesting a large part of proteins spontaneously localized on the liposome membrane (Supplementary Fig. 8). To visualize the membrane localization, PlsX and PlsY were synthesized inside GUV in the form of GFP-fusion proteins. We observed the synthesis of GFP-PlsX protein inside, but they did not localize on the GUV membrane (Fig. 2C, Supplementary Fig. 9). Since PlsX is a peripheral membrane protein that requires a negative charge on membrane surface for localization ^21, 22^, we added 1-palmitoyl-2-oleyl-sn-glycero-3-phospho-rac-(1-glycerol) (POPG) in the lipid composition of GUV in the range of 0-50 % (% mol.) to observe that effect on the membrane localization. We found a weak tendency of the membrane localization at 25 % POPG, but a large part of GFP-PlsX localized at 50 % POPG (Fig. 2C, Supplementary Fig. 9, S10). Interestingly, this movement became more significant when Ficoll was introduced inside the GUV. Ficoll is often used as a bulky polymer that brings a crowding effect in a vesicle or a water-in-oil emulsion ^23^. The effect of Ficoll on the membrane localization of peripheral proteins has been observed in the case of FtsZ ^5^. When Ficoll was added, the formation of fluorescent spots was observed even at 0% or 10% POPG (Fig. 2C, Supplementary Fig. 9), perhaps indicating non-ordered polymerization of GFP-PlsX. On the other hand, at 25% POPG, most proteins localized on the vesicle membrane that is quite different from that of the Ficoll minus. In 50% POPG, visually all proteins localized on the GUV membrane. These results showed that the rate of membrane localization of PlsX synthesized inside GUV is substantially enhanced by the negative charge on the membrane by POPG and the molecular crowding effect by Ficoll. In contrast to PlsX, PlsY localized to the GUV membrane by its hydrophobic interaction. We observed the membrane localization of PlsY-GFP which was synthesized in the presence of Ficall inside GUV containing 50% POPG (Fig. 2D, Supplementary Fig. 11).

**Figure 2.**
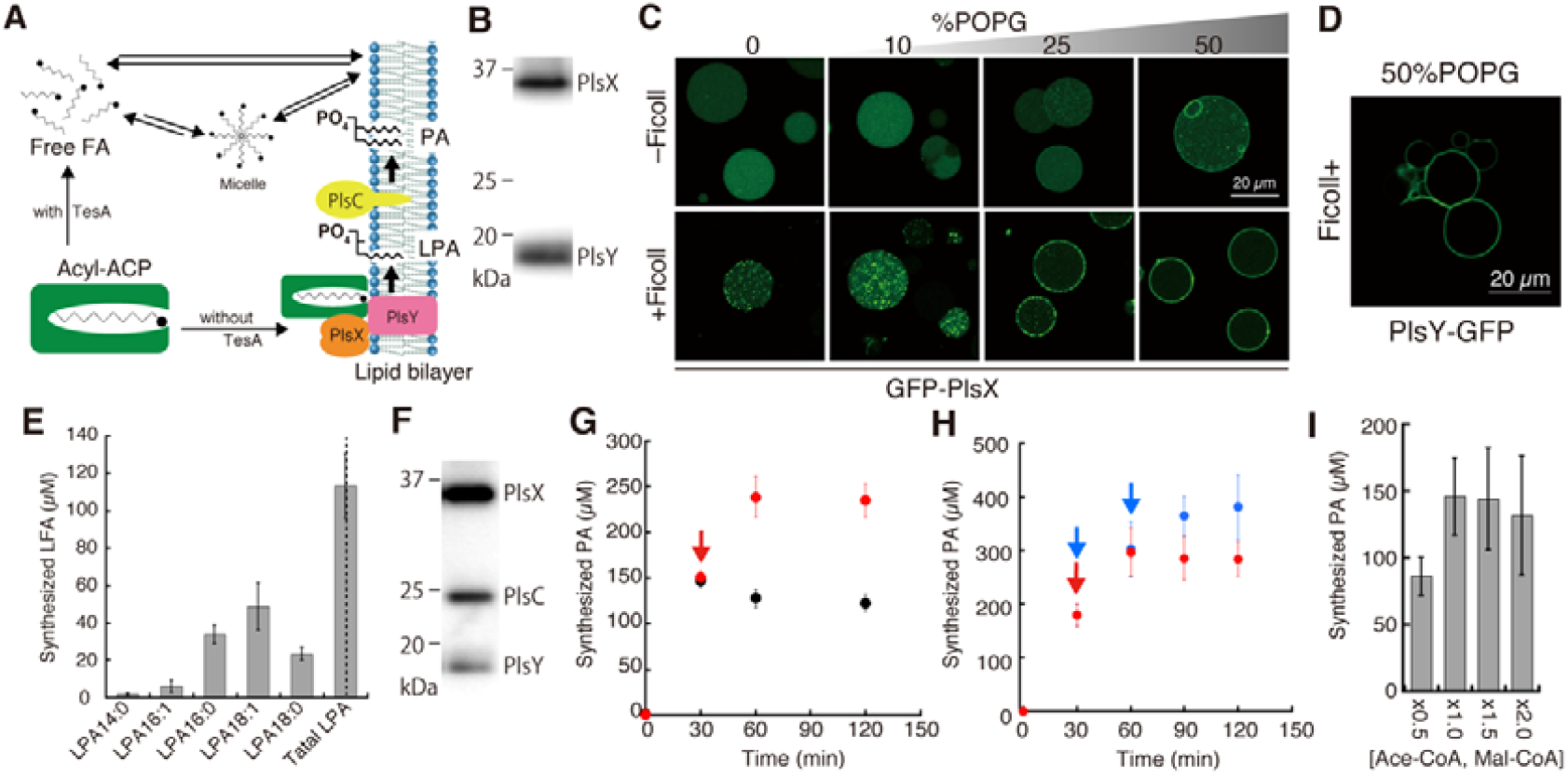
Phospholipid synthesis by cell-free synthesized acyltransferases. (A) Schematic of phospholipid synthesis pathway in the cell-free system. (B) SDS-PAGE image of cell-free synthesized PlsX and PlsY. Both proteins contain a six-histidine-tag at the C-terminus of the protein. The bands were visualized by anti-histag antibody. (C) Confocal microscopy image of the GFP-fusion PlsX synthesized inside giant unilamellar vesicles (GUVs) in the presence or absence of Ficoll. The percentage of POPG (%mol) in the lipid composition of GUV is described above the images. Scale bar: 20 µm. (D) Confocal microscopy image of GFP-fusion PlsY, as same as (C). (E) Synthesis of lysophosphatidic acids (LPAs) by PlsX and PlsY synthesized in a bulk cell-free system. (F) SDS-PAGE image of cell-free synthesized PlsX, PlsY, and PlsC, as same as (B). (G) Synthesis of phosphatidic acids (PAs) by cell-free synthesized PlsX, PlsC, and PlsC. At 30 minutes, the resources (acetyl-CoA, malonyl-CoA, and NAD(P)H) (red) or buffer (black) were additionally supplied into the reaction mixture. (H) Increase of PA synthesis by supplying the resources at 30 and 60 minutes (blue) or only 30 minutes (red), as well as (G), (I) Effect of the various concentrations of resources supplied from the beginning of PA synthesis reaction. The concentrations of ×1.0 [Ace-CoA, Mal-CoA] is 2 and 4mM, respectively. Error bars indicate the standard deviation of at least triplicate measurements. LPA: lysophosphatidic acid.

### Lysophospholipids synthesis by expressed PlsX and PlsY

To examine whether the cell-free synthesized PlsX and PlsY exhibit acyl-transfer activities in cooperation with each other, we synthesized the two enzymes in a bulk PURE system containing liposomes which consist of 50 % POPC and 50 % POPG (%mol). After the protein synthesis, we combined the PURE system with a Fab enzymes mixture in which acyl-ACP has been formed in advance through a single round of fatty acid synthesis without TesA. In the absence of TesA, fatty acid synthesis has been sufficiently suppressed at least for three hours of incubation (Supplementary Fig. 12). The resulting mixture allowed LPA synthesis following fatty acid synthesis with acetyl-CoA, malonyl-CoA, and NAD(P)H. The products were extracted and analyzed by an LCMS. The result showed that LPAs containing several types of the hydrocarbon chain were detected (Fig. 2E). This means that the synthesized PlsX and PlsY localized onto the liposome membrane and catalyzed LPA synthesis in collaboration with fatty acid synthesis. LPA18:1 was found as the most abundant product, followed by LPA16:0 and LPA18:0. Because only a small amount of the LPA16:1 product was detected, it seems that the synthesis of unsaturated fatty acids accomplished until the ultimate cycle, i.e. 18:1. In fatty acid synthesis alone, saturated fatty acid synthesis proceeded to C16:0, but C18:0 was barely synthesized (Fig. 1C, D, G, and H). On the other hand, when the lipid synthesis further advanced to LPA, a significant amount of LPA18:0 was produced that corresponds to 40 % of the total saturated LPA products (Fig. 2E). That may suggest that the saturated 16:0 acyl-ACP is sensitive to TesA, resulting in the release of 16:0. Contrary, when TesA was absent, acyl-ACP had much chance to elongate up to 18:0, thus increased the LPA18:0 product. The total amount of the synthesized LPAs was 120 µM. It means that about 65 % of the malonyl-CoA and 20 % of the acetyl-CoA converted into LPA (e.g. as LPA 18:0). Furthermore, since 15 µM ACP was presented to the system, the turnover of ACP is estimated to be eight cycles or more on average. We also found successful LPA synthesis in a one-pot reaction in which the Fab enzymes existed during the protein syntheses.

### Phosphatidic acid synthesis by expressing PlsX, PlsY and PlsC in the cell-free system

Although we detected successful LPAs synthesis, the yield (Fig. 2E) did not exceed that of fatty acid synthesis (Fig. 1C, D, G, and H). It may be due to the functional inhibition of PlsY by the synthesized LPA ^24^. To overcome this problem, we additionally introduced a gene for PlsC to continue the phospholipid synthesis one step further toward phosphatidic acid (PA) (Fig. 2A). When PlsX, PlsY and PlsC were co-synthesized (Fig. 2F), 150 µM PA was synthesized until 30 minutes (Fig. 2G). However, the yield did not increase after that. We hypothesized that the termination of PA synthesis was caused by the depletion of substrates and NAD(P)H. To test this, we next additionally supplied the resources (acetyl-CoA, malonyl-CoA, and NAD(P)H) at the point of 30 minutes of the reaction to the same concentration as the initial condition. The result showed that the amount of synthesized PA increased up to 250 µM at 60 minutes (Fig. 2G), whereas the yield kept constant or slightly decreased when we did not supply the additional resources. These results indicate the resources for PA synthesis depleted in the cell-free mixture within 30 minutes. We once again supplied the same resources at the 60 minutes, resulting in a further increase to 350-400 µM PA at 90-120 minutes (Fig. 2H). Here, note that one PA contains two fatty acid chains. Therefore, the obtained maximal yield is equivalent to 700-800 µM fatty acid or LPA these are single-chain lipids. This is a remarkable enhancement compared to so far synthesized fatty acids or LPAs. It is an order of magnitude higher than the previous studies reporting 30µM phospholipids synthesis using acyl-CoA as a hydrocarbon chain source ^13, 14^. Because the yield was still growing even after the second dose, we expect that the phospholipid synthesis would increase further by adding more resources. Contrary, when we added 1.5 or 2 times more resources at the beginning of the reaction, the increase of PA synthesis was not observed or even slightly decreased (Fig. 2I). These results indicate that excess substrates adversely affect lipid synthesis. Based on the above results, we conclude that continuous reaction flax from fatty acid synthesis to phospholipid synthesis is highly important for maximizing the yield of lipid synthesis, avoiding the accumulation of synthetic intermediates and reaction suppression by their negative feedback on the lipid synthesis.

### Phosphatidic acid synthesis from acetic acid and bicarbonate

The constructed cell-free phospholipid synthesis system uses acetyl-CoA and malonyl-CoA as hydrocarbon sources. This system produces, at the same time, by-product CoA as the reaction progresses. For example, 18 CoAs will be produced per one PA (e.g. 18:0 or 18:1) synthesis. This will be a problem when producing 1 mM phospholipid inside a GUV, as it will accumulate a considerable amount of CoA. To avoid this problem and make the system more sustainable, we redesigned the system that recycles CoA for producing acetyl-CoA and malonyl-CoA. We further purified acetyl-CoA synthetase (ACS) and acetyl-CoA carboxylase (Acc) ABCD (Fig. 3A). ACS synthesizes acetyl-CoA from acetic acid and CoA using the energy of ATP, and AccABCD synthesizes malonyl-CoA from acetyl-CoA and bicarbonate. We added these enzymes during the cell-free syntheses of PlsX, PlsY, and PlsC. Then, CoA, NAD(P)H, and KHCO_3_ were supplied to start the PA synthesis. We did not add acetic acid because the PURE system is containing it enough as a counter ion of the magnesium ^2^. The PURE system also contains ATP and an energy regeneration system ^2^. The result showed that about 80 µM PA were synthesized in 60 minutes reaction (Fig. 3B). No PA was synthesized in the absence of CoA, ACS, AccDA, or AccBC (Supplementary Fig. 13). These results support that acetyl-CoA and malonyl-CoA were synthesized from CoA and low-molecular compounds (acetic acid and bicarbonate) and, continuously, consumed by the cell-free synthesized enzymes to synthesize PA (Supplementary Fig. 14). This system has an advantage especially for synthesizing phospholipids in the enclosed space of GUVs.

**Figure 3.**
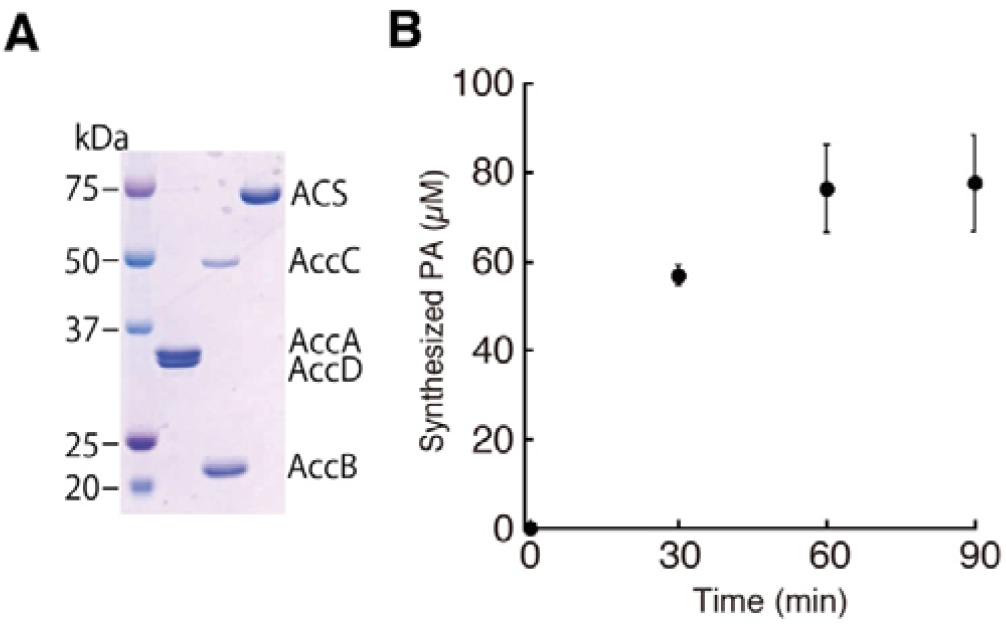
Phosphatidic acid synthesis by synthesizing acetyl-CoA and malonyl-CoA. (A) An SDS-PAGE image of the purified acetyl-CoA carboxylase DA (AccDA), acetyl-CoA carboxylase BC (AccBC), and acetyl-CoA synthase (ACS). The positions of molecular size are described on the left side of the gel image. (B) Phosphatidic acid synthesis by synthesizing acetyl-CoA and malonyl-CoA by ACS and AccABCD. The reaction was initiated by adding CoA. Acetic acid and potassium bicarbonate were used as carbon sources.

### Expression of pathogenic genes for phospholipid synthesis

In this system, phospholipids are synthesized *in vitro* through fatty acid synthesis and gene expression of acyltransferases that are membrane protein enzymes. The system can evaluate the function of the gene products within 24 hours, so we think it is applicable for other bacterial acyltransferases, such as pathogenic bacteria. To prove this, we tried to synthesize *plsX, plsY*, and *plsC* genes of methicillin-resistance *Staphylococcus aureus* (MRSA) and JCVI-Syn3.0, which is a designed synthetic genome based on *Mycoplasma mycoides* ^25^. All proteins were synthesized in the PURE system supplemented with liposomes which consists of 50 % POPC and 50 % POPG (%mol). We found that the most synthesized proteins were found in the supernatant after centrifugation (Supplementary Fig. 15), suggesting localized onto the liposome membrane. Only MRSA PlsY (54s% sol.) and MRSA PlsC (5 % sol.) showed somehow low membrane localization.

Next, we performed PA synthesis by coupling fatty acid synthesis and protein synthesis of PlsX, PlsY, and PlsC of MRSA or Syn3.0. The result showed that PAs were synthesized when MRSA genes were expressed (Fig. 4A). The total yield was a little bit higher than that of the *E. coli* case. The pattern of the acyl-chain variety in the PA products was different from that of the *E. coli* case (Fig. 4B, Supplementary Fig. 16), i.e. the proportion of short acyl-chains including C14:1 and C16:1(0) tends to be relatively higher than the long chains compared with the *E. coli* case. On the other hand, no significant PA synthesis was detected when the Syn3.0 genes were expressed. To find out the reason, we compared alignment scores of each gene in the *E. coli* vs MRSA and the *E. coli* vs Syn3.0. However, a remarkable difference was not found between them (Supplementary Text2). Next, we compared the alignment score of ACP that directly interacts with PlsX in the cytosol. We found that the amino acid sequence of MRSA ACP showed high homology (Score 57.14) with that of *E. coli* ACP (Supplementary Text2). Contrary, the amino acid sequence of Syn3.0 ACP did not show significant homology (Score 24.66). These results imply that the synthesized Syn3.0 PlsX could not crosstalk to *E. coli* ACP, which presents in the fatty acid synthesis system, so PA synthesis did not occur. On the other hand, MRSA PlsX was able to crosstalk with *E. coli* ACP, resulting in the synthesis of PA.

**Figure 4.**
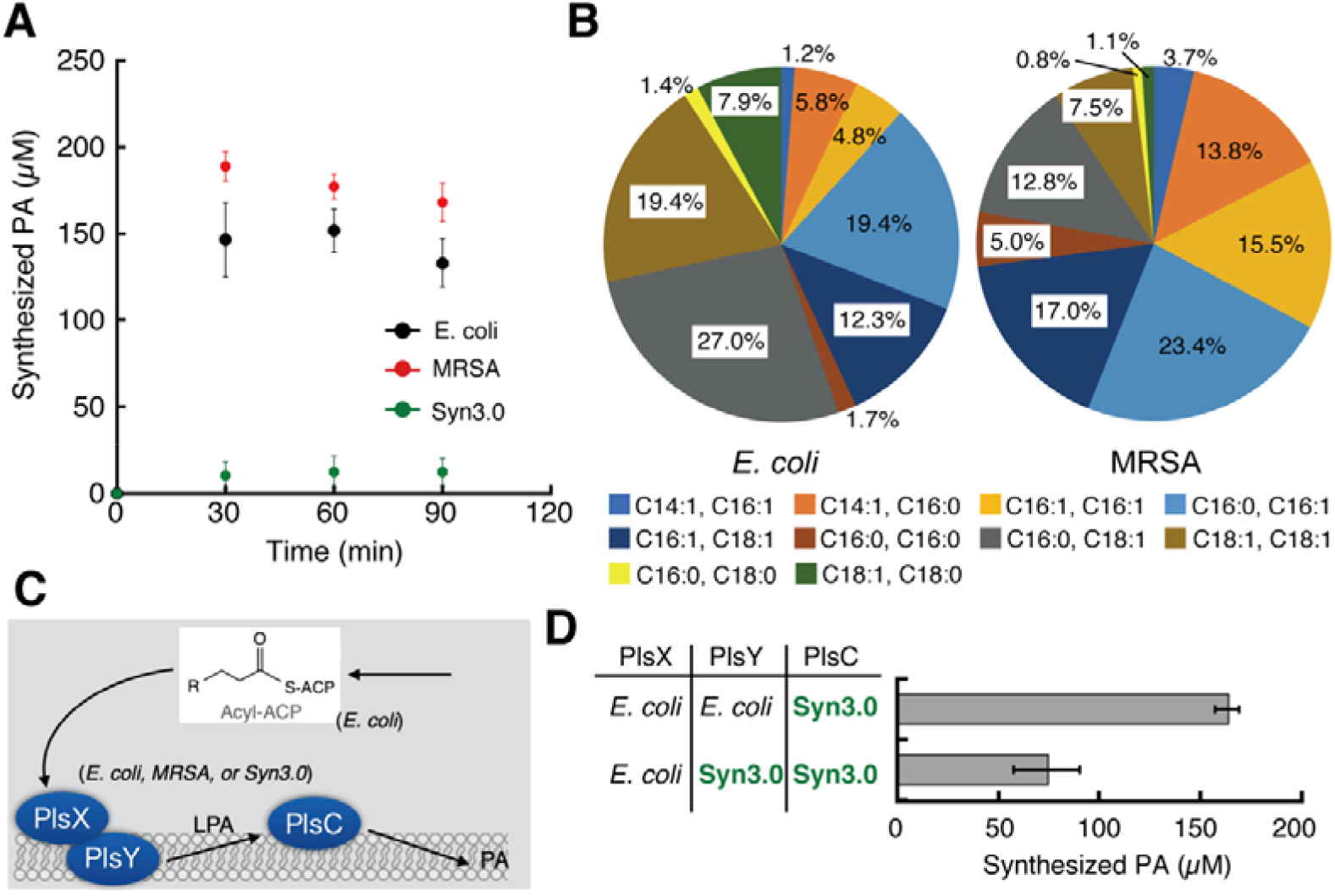
Cell-free expression of pathogenic bacterial acyltransferase genes. (A) Phosphatidic acid synthesis by cell-free synthesized PlsX, PlsY, and PlsC of *E. coli*, MRSA, or Syn3.0. (B) The proportion of acyl-chains in the synthesized phosphatidic acid from *E. coli* and MRSA genes. The fatty acid composition at 30 minutes in (A) was compared. (C) Schematic of phosphatidic acid synthesis pathway from Acyl-ACP. (D) Phosphatidic acid synthesis by cell-free synthesized *E. coli* PlsX&PlsY and Syn3.0 PlsC or *E. coli* PlsX and Syn3.0 PlsY&PlsC. Error bars indicate the standard deviation of at least triplicate measurements.

If this is true, PA could be synthesized in the set of *E. coli* PlsX&PlsY and Syn3.0 PlsC, because PlsX&PlsY and PlsC are not directly interacting on a cell membrane (Fig. 4C). The result showed PA synthesis was detected when *E. coli* PlsX&PlsY and Syn3.0 PlsC were expressed (Fig. 4D), supporting our hypothesis that is correct. We also found the PA synthesis when *E. coli* PlsX and Syn3.0 PlsY&PlsC were expressed, but the synthesis efficiency was low (Fig. 4D).

Since there is no homologous between the bacterial and human acyltransferases, attacking these functions may inhibit the growth of pathogenic bacteria such as MRSA and mycoplasma. We believe that our cell-free system is valuable as an in vitro system that allows easy and quick screening of antibacterial chemicals targeting a root of phospholipid synthesis pathway.

## Conclusions

In this study, we constructed the cell-free phospholipid synthesis system by assembling three cell machinery, fatty acid synthesis, protein synthesis, and phospholipid synthesis. Additionally, the acetyl-CoA and malonyl-CoA synthesis from CoA were also constituted and implemented in the system. These have been constructed from the individually isolated components so that we can comprehend all contents in the cell-free system and their concentrations. This fact is crucial for building a living cell and, especially, for defining essential small molecular compounds that cannot be captured from genetic information. By doing the construction, we have found that the reaction rate of bacterial phospholipid synthesis is much faster than other similarly complicated systems, such as protein synthesis, despite being a complex system of multi-components. We also found that lipid synthesis is a risky process for the vital activity of cells, i.e. synthesizing hydrophobic molecules in an aqueous cytosol space brings serious risk for the enzyme activities. To avoid such risk, cells synthesize a nascent fatty acid within the cradle of ACP and directly deliver it to the cell membrane via PlsX and PlsY. This is very rational also in terms of efficiently growing a cell membrane. Even after converted fatty acids into LPAs, there is another risk at the level of the lipid membrane, i.e. LPAs, which have inverted-cone shapes, can form a pore on the membrane. Cells counteract this problem by rapidly converting LPAs to phospholipids that can maintain a formation of a stable lipid bilayer. In summary, phospholipid synthesis is a very risky process for cells, but cells avoid it by an entirely rational mechanism.

Our new cell-free system has achieved phospholipid synthesis in higher concentrations than the previous schemes. The method of our previous work in 2009 used the same cell-free system (PURE system) for the synthesis of acyltransferases (plsB and PlsC). But only a limited number of phospholipids were synthesized because of the low productivity of the early version of the PURE system. Danelon’s group methods in 2016 and 2020 used a more productive PURE system but did not provide sufficient phospholipid synthesis to deform the GUV membrane. In both cases, acyl-CoA was employed as a source of hydrocarbon chains for phospholipid synthesis, and we believe that it is the cause of the low phospholipid productivity. Unlike these approaches, we reconstructed the whole process of hydrocarbon chain synthesis and combined it with phospholipid synthesis in a one-pot. By developing such fluxes, we could avoid the accumulation of fatty acids and LPAs and skip the inhibitory process by them. As a result, we were able to increase the yield of phospholipid synthesis to an order of magnitude higher than before. We showed that the reaction continued until depleting the substrates and electron donors (Fig. 2G and H). We are currently investigating the system to produce more phospholipids. Perhaps the productivity may increase if the reaction goes beyond the synthesis of negatively charged PAs to the synthesis of the zwitterionic triglycerides phosphatidylcholine or phosphatidylethanolamine.

The next challenge of this work is to encapsulate inside GUV and to perform phospholipid synthesis therein. Unlike the bulk reaction, free diffusion of the molecules is restricted inside GUV. So, it is necessary to consider the membrane permeability of the molecules required for internal phospholipid synthesis. Additionally, it should be considered that the vesicle membrane exhibits morphological changes only in lipid dynamics, even if phospholipids are not being synthesized internally. To determine that the vesicle deformation is exactly occurred due to the internal reaction, confirming a sufficient phospholipid amount is synthesized in the bulk condition is first essential.

Lastly, we examined the genes of acyltransferases from pathogenic bacteria. The synthesized pathogenic PlsX exhibited its acyltransferase activity in the cell-free system when its host ACP is highly homologous to the *E. coli* ACP. Even the homology is low, the synthesized pathogenic PlsY and PlsC can co-work with *E. coli* PlsX and PlsY. This fact indicates that our cell-free system is available as an *in vitro* evaluation system for the acyltransferases of interest without awkward membrane protein purification. Such a system is also applicable for exploring candidate drugs or antibodies which attack the early step of phospholipid synthesis of multiple-drug-resistant bacteria or nosocomial infection-causing bacteria.

The developed cell-free system in this study is designed for synthesizing phospholipid inside GUVs that eventually toward the construction of a self-propagating artificial cell. To accomplish this, mM scale phospholipids must be produced by the internal metabolic reaction. We believe that our result is a big step toward solving this problem and promising the construction of self-growing and self-dividing artificial cells.

## Materials and methods

*Materials*—1-palmitoyl-2-oleoyl-glycero-3-phosphocholine (POPC),

1-palmitoyl-2-oleoyl-sn-glycero-3-phospho-(1’-rac-glycerol) (POPG),

1-myristoyl-2-hydroxy-sn-glycero-3-phosphate (14:0 Lyso PA),

1-palmitoyl-2-hydroxy-sn-glycero-3-phosphate (16:0 Lyso PA),

1-stearoyl-2-hydroxy-sn-glycero-3-phosphate (18:0 Lyso PA),

1-oleoyl-2-hydroxy-sn-glycero-3-phosphate (18:1 Lyso PA),

1,2-dipalmitoyl-sn-glycero-3-phosphate (DPPA),

1-palmitoyl-2-oleoyl-sn-glycero-3-phosphate (POPA), and

1,2-dioleoyl-sn-glycero-3-phosphate (DOPA) were purchased from Avanti Polar lipids. Fatty acid LC/MS mixture (#17942) was purchased from Funakoshi (Japan).

^13^C-Acetyl-CoA, ^13^C-Malonyl-CoA, Acetyl-CoA, Malonyl-CoA, NADH, NADPH, and *sn*-glycerol-3-phosphate were purchased from Sigma Aldrich. PURE*frex* 2.0 and its Sol.I buffer, which was customized for volume down, and DnaKJE mix were given or purchased from GeneFrontier (Japan).

### Overexpression and protein purification

All Fab enzymes, ACP and TesA were overexpressed and purified as previously reported by Yu et al. (Yu 2011 *PNAS*) with some modifications (Supplementary Table2). Briefly, for FabA/B/D/F/G/H/I/TesA, the transformed cells were cultivated in 1 L LB medium supplied 50 µg/mL kanamycin at 37 °C. At OD_600_ 0.6-0.8, 0.1 mM Isopropyl-β-D-thiogalactoside (IPTG) was introduced for induction, and the cultivations were continued for 3 hours. The cells were then collected by centrifugation 5,500 × g for 20 min at 4 °C, washed with Buffer A (50 mM Tris-HCl (7.6), 2 mM dithiothreitol (DTT), 10 % glycerol), and stored at −80 °C until use. The frozen cells were dissolved in 35 mL Buffer A on ice and disrupted by sonication (5 sec on, 3 sec off power 40%, total 5 min). The unbroken cells and debris were removed by centrifuging 30 k × g for 1 hour at 4°C. The resulting supernatants were collected and injected to AKTA™□ pure equipped HisTrap HP column (GE Healthcare), which was pre-equilibrated with Buffer A. The HisTagged proteins were eluted by a gradient of Buffer B (Buffer A with 1M Imidazole) from 3% to 50%B. The target proteins were collected and desalted using Amicon (10kDa cut) (Millipore) before subjecting to the HiTrap Q column (GE Healthcare). The injected proteins were washed and eluted by a gradient of Buffer C (Buffer A with 1M NaCl) from 0 % to 100 %B. The target proteins were collected and desalted by Amicon. Finally, concentrated proteins were determined those concentrations with Bradford (Takara Bio) or BCA (Thermo Fisher Scientific) protein assay kit, and stored at −80 °C. In general, the purified proteins are stable for f at least a few months.

ACP was co-expressed with a 4’-phosphopantetheinyl transferase (SFP) because ACP needs post-translational modification by SFP to form Holo-type ACP (Holo-ACP). The source of the *sfp* gene is from *bacillus subtilis*, and the codon was optimized for *E. coli* expression system when we ordered it as a synthetic gene (Supplementary Table3). The gene was inserted between NcoI and BamHI sites of pCDF-1b (Novagen) vector using the primers sfp-fw and sfp-rv (Supplementary Table4), resulting in pCDF-sfp (Supplementary Table2). The *E. coli* BL21 (DE3) cells harboring pTL1415 which carrying the *acp* gene and pCDF-sfp were cultivated in TB (Terrific Broth) medium supplied kanamycin and streptomycin until OD600 arrived at 0.6. After the induction with 1mM IPTG, cells were collected at OD600 1.9 and washed with 50 mM Hepes-KOH (pH7.6). The obtained cell pellets were dissolved in Buffer A and disrupted by French Press at 500 bar (7,000 psi) three times. After removing cells unbroken and debris by centrifugation, the supernatant was subjected HisTrap HP column as in the case of the Fab proteins, using the Buffer A and B containing 300 mM NaCl. The target proteins were collected and desalted for subjecting MonoQ 4.6/100PE column (GE Healthcare) using Buffer A and C as shown in Fab proteins. The collected target proteins were desalted by nap-5 and concentrated by Amicon. Concentrations of the protein were determined by Protein Assay (BioRad). MALDI-TOF-MS was used to confirm the Holo-type of the purified ACP.

Because FabZ tends to aggregate easily, we modified the expression and purification method as follows. The expression plasmid pXY-FabZ was introduced to *E*. *coli* C43(DE3) strain cells. The cells were cultivated at 37 °C in 1 L LB medium supplied kanamycin until the A600 0.6, then, 1mM IPTG was added followed further 2 hours cultivation. The resulting cells were collected and washed with 50 mM Hepes-HOH (pH 7.6), then stored at −80 °C until use. The cell pellets were resuspended in Buffer D (100 mM potassium phosphate (pH 7.4), 300 mM NaCl, 1mM DTT) and disrupted by French Press as same as ACP. After removing cells unbroken and debris, the supernatants were subjected HisTrap HP column with Buffer D and E (Buffer D with 1 M Imidazole). The FabZ were eluted with a gradient of Buffer E from 10% to 100%. The fractions containing FabZ were collected and desalted by repeating dilution with Buffer D and concentrating by Amicon. Note that the purified FabZ should not be concentrated to a high concentration to prevent aggregation. We did not proceed to further purification because FabZ is unstable, and only HisTrap purification is enough for purity. If it was difficult to determine the FabZ concentration by the Protein Assay, the concentration was defined by the band intensity on SDS-PAGE analysis comparing to the standard protein whose concentration had already been determined. Interestingly, the aggregated FabZ was dissolved by mixing with ACP. This might be due to the complex formation of FabZ and ACP that has been reported by Zhang *et al*.26. Therefore, FabZ should be stored as a mixture with ACP in the molar ratio of 1:30 (FabZ: ACP).

Acetyl-CoA synthetase (ACS) was overexpressed from the synthetic gene (Supplementary Table3) located between NdeI and XhoI sites in the pET-28a vector harboured in *E. coli* C43(DE3) cells, referring to a previous report^27^. Cells were cultivated at 37 °C in LB medium supplied kanamycin until the A_600_ 0.4, then, 0.4 mM IPTG was added followed by further three hours cultivation. The resulting cells were collected and washed with 100 mM KPO_4_ (pH 7.4), then stored at −80 °C until use. The frozen cell pellet was dissolved in Buffer D and disrupted by French Press as in the case of ACP. After removing cells unbroken and debris, the supernatants were subjected by AKTA Avant 25 (GE Healthcare) equipped HisTrap HP column with Buffer D and E. The ACS was eluted by the gradient of Buffer E from 10% to 50% for 20 min by flow rate 2.5 ml/min. The fractions containing ACS were collected and desalted by repeating dilution with Buffer D and concentrating by Amicon 30K Da cut off. The resulting sample was further purified by MonoQ 4.6/100PE column (GE Healthcare) using Buffer F (50 mM Tris-HCl (pH 8.0), 10 % glycerol) and Buffer G (Buffer F with 1M NaCl). ACS was eluted by the gradient of Buffer G from 0% to 100% for 20 min by flow rate 1.0 ml/min. The collected ACS was desalted and concentrated using Amicon. We did not cut off HisTag because the protein aggregates.

AccBC complex was purified as previous reports28, 29, with slight modifications. The polycistronic genes for AccB and AccC were obtained as a synthetic gene (Supplementary Table3) with codon optimization for *E. coli* expression system (FASMAC), which inserted between NdeI and XhoI sites of the pET28a vector, resulted in pET28a-accBC. In this construct, a six-histidine tag was inserted at the N-terminus of the accB gene. The gene of birA, biotin-[acetyl-CoA-carboxylase] ligase, is also codon-optimized and inserted in the pCDF-1b vector using NcoI and XhoI sites, resulted in pCDF-birA. Both pET28a-accBC and pCDF-birA were co-introduced into C43 (DE3) *E. coli* cells (Lucigen) and then cultivated at 37°C in LB medium supplemented with kanamycin and streptomycin. At the OD660 0.4, 0.5 mM IPTG and 2µM biotin were added, then, followed by further cultivation for four hours. Cells were collected and dissolved in Buffer H (20 mM potassium phosphate (ph8.0), 500 mM NaCl, 2-mercaptoethanol, protease inhibitor cocktail), then disrupted by French Press as described above. After removing cells unbroken and debris, the lysate was subjected by the HisTrap HP column with Buffer H and I (Buffer H with 1 M imidazole). AccBC protein was eluted by the gradient of Buffer I from 2% to 100% for 20min by flow rate 5.0 ml/min. The fractions of AccBC were collected and desalted using Amicon (10KDa cut off) and Buffer J (50 mM Tris-HCl (8.0), 100 mM NaCl, 10 % glycerol). The desalted proteins were further purified by the MonoQ column with Buffer J and K (Buffer J with 1 M NaCl). AccBC were eluted by the gradient of Buffer K from 0% to 100% for 20min by flow rate 1.5 ml/min. The collected AccBC fractions were buffer exchanged for Buffer L (20 mM Tris-HCl (8.0), 300 mM NaCl, 10% glycerol) while being concentrated by Amicon (10KDa cut off). The concentration was determined by the BCA protein assay kit. Finally, the efficiency of biotinylation of the purified AccB biotin carboxyl carrier protein of acetyl-CoA carboxylase) was confirmed. 1 µg AccBC sample was dissolved in 25.5 µL 50 mM ammonium bicarbonate (pH 8.0), then 1.5 µL fresh 500 mM DTT was added. After boiling at 95 °C for five minutes, 3 µL 500 mM iodoacetamide was added, followed by a 20 min incubation at room temperature. For digestion of the protein, 0.5 µL 5ng/µL Gluc-C (Promega) was added and incubated for three hours at 37 °C, followed additional supplying 0.5 uL Gluc-C and incubation at 30 °C for overnight. The digestion reaction was terminated by the addition of 3 µL of 50 % trifluoroacetic acid (TFA). The sample was then purified by a C18 spin column (Thermo Fisher) following the manufacture’s protocol. The eluted sample was dried up and dissolved in 5 % acetonitrile solution supplied 0.5 % TFA, before analyzing by Nano-LCMSMS (Thermo Fisher). From the obtained data, the area intensity of the biotinylated fragment (AMkMMNQIE, *m*/*z*=661.29232, *z*=2, where k=biotinylated lysine residue) was compared with that of a non-biotinylated fragment (AMKMMNQIE, *m*/*z*=548.25360, z=2). Generally, more than 50 % of the purified AccB was biotinylated.

AccDA was expressed from pACS27530 introduced in C43 (DE3) *E. coli* cells. pACS275 contains the genes of AccD, where a six His-tag sequence was added at the C-terminus, and AccA. The protein purification was carried out following a previous report30, with slight modifications. The cells were cultivated at 37 °C until the OD660 reached 0.5. 0.5 mM IPTG was added then cells were further cultivated at 25 °C for 4 hours. Cells were collected and washed with Buffer M (20 mM Tris-HCL (8.0), 500 mM NaCl, 10 % glycerol). For disruption of the cells, 2 µL 250 U/µL Benzonase (Merck) and 20 mM lysozyme were added into a 40 mL cell solution and kept on ice for one hour. After three times freeze-and-thaw, unbroken cells and debris were removed by centrifugation and the supernatant was passed a 0.45 µm filter. The resulting lysate was supplemented 50 mM (final concentration) imidazole, then subjected by the HisTrap HP column with Buffer M and N (Buffer M with 1 M imidazole). The sample was washed with 5 % Buffer N for 10 column volume by flow rate 5.0 mL/min, then eluted by the gradient of Buffer N from 5 % to 25 % for 20 column volume. AccDA fractions were collected and diluted five times with Buffer O (20 mM Tris-HCL (8.0) and 10 % glycerol). The resulting sample was loaded to the HiTrap Capto Q column (Cytiva) and washed 10 % Buffer P (Buffer O with 1 M NaCl) for 10 column volume. AccDA sample was eluted by the gradient of Buffer P from 10 % to 100 % for 20 column volume. The collected fractions were concentrated and buffer exchanged with Buffer L using Amicon (10 KDa cut off).

The compositions of all buffers used for protein purifications are shown in Supplementary Table5.

### Liposome preparation

40 mg POPC powder or a mixture of 20 mg POPC and 20 mg POPG powder was dissolved in chloroform within a round-bottom 50 mL flask and briefly processed by bath sonication. Then chloroform was removed by rotor evaporator and placed under low pressure within a desiccator for overnight to remove the residual chloroform completely. The resulting lipid film was dissolved with 1 mL 50 mM Hepes-KOH (pH 7.6) by vortexing and sonicating. The lipid solution was next processed by micro extruder using 1 µm and 200 nm pose size polycarbonate membrane in a stepwise fashion. The prepared liposome solution was divided into micro tubes and stored in −80 °C. The liposome sample is generally stable for at least years, but it should be passed a bath sonication before use.

### In vitro fatty acid synthesis

*In vitro* fatty acid synthesis reaction was prepared as described in Supplementary Table6, based on the report by Yu *et al*. (Yu 2011 *PNAS*). ^13^C-stable isotope-labeled acetyl-CoA and malonyl-CoA were used as substrates to distinguish the products from surrounding fatty acids. The reaction was generally carried out at 30 °C for 30 min. The resulting reaction mixtures were diluted to 20-fold with 50 % MeOH and centrifuged at 20,000 × g for 30 min at 16 °C. The resulting supernatants were collected and placed in glass vials. Synthesized fatty acids were analyzed by Shimadzu LCMS-2020 system equipped with InertSustain^⍰^ Phenylhexyl column (3 µm, 2.1 mm Cat. No. 5020-89128, GL Science, Japan) with a guard column (GL Science, Japan). Fatty acids were separated in 60 % B isocratic elution mode using mobile phase A (20 mM ammonium acetate) and B (100 % acetonitrile). The column oven was 35 °C, and the flow rate was 0.14 mL/min. Elution was analyzed by selected ion monitoring (SIM) mode monitoring m/z of 211.9 (C^13^12:0), 241.2 (C^13^14:0), 269.2 (C^13^16:1), 271.2 (C^13^16:0), 299.2 (C^13^18:1), and 301.3 (C^13^18:0) (Supplementary Table7) in the negative-ion mode. For the quantification, the area counts obtained from the peaks were converted into fatty acid amounts using the standard curve made from the analysis of the commercially available standard fatty acid mixture with defined concentrations. The standard fatty acids samples were measured each time of the experiment. An example of the standard curve is shown in Supplementary Fig. 17.

### Template DNA preparation for cell-free protein synthesis

The genes expressed in the PURE system were summarized in Supplementary Table8. All genes were prepared as a plasmid DNA by InFusion cloning or a linear DNA by PCR (Supplementary Table9) using the designed primer sets (Supplementary Fig. 4) and template DNAs (Supplementary Fig. 3). The *E. coli* gene of *plsX* and *plsY* was obtained from K12 strain cells and inserted into the pET28a vector. Using the resulting plasmids, we introduced a gene for super-folder green fluorescence protein (*sfgfp*)^6^ to the upstream of *plsX* or the downstream of *plsY* by InFusion to generate a GFP-PlsX_wt_ or PlsY_wt_-GFP plasmid, respectively (see Supplementary Table8 #7&8). These were used for expression in GUV.

Because the productivity of PlsX in the PURE system was not so high, we reduced the GC-contents just after the initial codon by introducing silent mutations by PCR (see Supplementary Table8 #1). We used this liner DNA for cell-free LPA or PA synthesis. In the same manner, we also prepared 6His-tagged *plsX* for the analysis of western-blotting, using an anti-HisTag antibody (see Supplementary Table8 #4). The *E. coli* gene of *plsY* for cell-free LPA or PA synthesis was obtained as a synthetic gene with codon optimization for the *E. coli* expression system (see Supplementary Table8 #2). As same as *plsX*, a 6His-tagged *plsY* was prepared by PCR (see Supplementary Table8 #5). The *E. coli* gene of *plsC* was obtained as a previously prepared plasmid pBT302_plsC^12^ (see Supplementary Table8 #3). This was used for cell-free PA synthesis and for preparing a 6His-tagged *plsC* DNA (see Supplementary Table8 #6). The *plsX, plsY*, and *plsC* genes of MRSA or Syn3.0 were obtained as synthetic genes with codon optimization for the *E. coli* expression system (see Supplementary Table8 #9-14). All these were used for cell-free PA synthesis and for preparing 6His-tagged DNAs (see Supplementary Table8 #15-20).

### Cell-Free Protein synthesis

Cell-Free gene expressions were performed using PURE*frex* 2.0 (GeneFrontier, Japan) according to the manufacture’s protocol. In general, linear template DNA(s) of *plsX, plsY*, and *plsC* were introduced in the reaction mixture supplied liposomes, sucrose, and DnaKJE chaperon mixture (DnaKJE mix), which is highly affect the solubility of PlsX^20^ (eSol: http://www.tanpaku.org/tp-esol/index.php?lang=ja) (Supplementary Table10). For the gene expression to LPA or PA synthesis, the custom Sol. I, which was adjusted to reduce the volume to 6 µL by GeneFrontier company, was used to making an space to add the additional components, such as glycerol-3 phosphate (G3P) and fatty acid enzymes mixture (FA mixture). The FA mixture was prepared as shown in Supplementary Table10, of which 2 µL was used to make 20 µL of the reaction mixture. For the gene expression to synthesize PA in the CoA recycling system, ACS, AccBC, and AccDA were additionally introduced using the water space of the mixture. In all cases, gene expression reactions were carried out at 37 °C for 2 hours using a thermal cycler.

### Solubility assay

The ratio of membrane localization of the synthesized PlsX and PlsY were assessed as a previous report^31^, using the *plsX*-6His and *plsY*-6His genes. Proteins were synthesized in the presence or absence of liposomes consist of 50% POPC and 50% POPG. In the case of MRSA and Syn3.0 proteins, the reaction mixture was centrifuged for 30 min at 20,000 × g at 4°C to separate liposomes in the supernatant from the aggregated products that appeared in the precipitation. In both cases, the proteins were analyzed by western blotting using an anti HisTag monoclonal antibody-conjugating horseradish peroxidase.

### In vitro phospholipid synthesis

After the gene expression, the reaction mixture was suplemented with NADH, NADPH, Acetyl-CoA, Malonyl-CoA, and potassium phosphate buffer as described in Supplementary Table10. For LPA or PA quantification by LCMS, we used [^13^C]-labelled Acetyl-CoA and Malonyl-CoA. LPA or PA synthesis was performed at 30 °C. The resulting mixture was pre-treated as same as a previous report^13^ before analyzing by LCMS. Briefly, 5 µL of sample synthesizing LPA or PA was diluted with 45 µL or 495 µL 100 % MeOH solution including 2 mM acetylacetone, respectively. The diluted sample was sonicated for 10 min and then centrifuged at 16,000 × g for 5 min at 15 °C. The supernatant was passed a 0.2 µm filter (Millex®-LG) and injected into LCMS2020 (Shimadzu). 0.025 ng/µL of DPPA, POPA and DOPA were added as an internal standard in the MeOH solution to assess lipid extraction efficiency.

### LCMS analysis for synthesized phospholipids

For separation of synthesized LPAs and PAs, a metal-free column L-column2 or L-column3 (CERI, JAPAN) were used to avoid adsorption of phospholipids on the metal surface of the column. The LCMS analysis method is based on Blanken *et al*.^13^ with slight modifications. Mobile phase A (water with 0.05% ammonium hydroxide and 2 mM acetylacetone) and B (80 % 2-propanol, 20 % acetonitrile, 0.05 % ammonium hydroxide, and 2 mM acetylacetone) were used for the isolation of LPA and PA. For LPA, column oven at 55 °C, flow rate 0.2 mL/min, and %B 30 were applied. 5 µL samples were injected, then eluted by a gradient from 30% to 100% B in 30 min followed. Elution was directly sprayed into the MS to analyze the products in negative mode. The m/z values of 395.0 (LPA^13^C14:0), 393.2 (LPA^13^C14:1), 425.1 (LPA^13^C16:0), 423.1 (LPA^13^C16:1), 455.1 (LPA^13^C18:0), and 453.1 (LPA^13^C18:1) were monitored as ^13^C-labeled products in SIM mode as shown in Supplementary Table7. For PAs, when L-colum2 was used, the column oven at 60 °C, the flow rate 0.2 mL/min, and %B 50 was applied. When L-column3 was used, the column oven at 45 °C, the flow rate 0.15 mL/min, and %B 50 was applied. In both cases, 5 µL samples were injected, then eluted by a gradient from 50% B to 100% B in 30 min followed. The m/z values of 679.5 (PA^13^C16:0/16:0), 677.5 (LPA^13^C16:0/16:1), 709.5 (LPA^13^C16:0/18:0), 707.5 (LPA^13^C16:0/18:1), 675.5 (LPA^13^C16:1/16:1), 705.5 (LPA^13^C16:1/18:1), 739.5 (LPA^13^C18:0/18:0), 737.5 (LPA^13^C18:0/18:1), and 735.5 (LPA^13^C18:1/18:1) were monitored as ^13^C-labeled products in negative SIM mode as shown in Supplementary Table7.

### Formation of giant unilamellar vesicle

encapsulation of the cell-free system inside GUV was performed as described in Berhanu et al. ^6^ with slight modifications. A 100 µL of 100 mM lipids dissolved in chloroform was added into 1 mL mineral oil within a grass vial containing a micromagnetic stirrer bar. The lipid oil mixture was gently stirred at 60 °C for 60-90 min to evaporate chloroform. After chilling the oil, 20 µL cell-free reaction mixture was added into 500 µL lipid oil solution, then vigorously vortexed for 1 min. The resulting lipid solution was overlaid onto a 500 µL outer solution in a 1.5mL micro test tube. The outer solution is the same composition as Sol. I of PURE*frex*2.0 omitting tRNA mixture, creatine phosphate, NTPs, but supplied 200 mM glucose. Next, the sample was subjected to centrifugation at 10,000 × g for 30 min then the upper oil phase was removed by pipetting. The formed GUV at the bottom of the test tube was collected carefully by taking 30 µL by a pipette. The collected GUV solution was supplied ATP as final concentration 1mM and incubated for 2 hours at 37 °C. After the reaction, GUVs were observed by confocal microscopy.

### Confocal microscopy observation

GUVs were observed by Nikon confocal microscopy system A1R using 488/509 (ex/em) wavelength for GFP observation. From the obtained image, line plot analyses were performed by NIS-Element software for measuring the fluorescent intensity of GUVs.

## Acknowledge

The expression plasmids for Fab enzymes, ACP, TesA were kindly supplied from Prof. Chaitan Khosla (Stanford Univ.). The expression plasmids for AccDA was kindly supplied from Prof. John E. Cronan (Univ. of Illinois). Dr Takashi Kanamori (GeneFrontier) kindly supplied the PURE system.

We thank Yoshi Kawai (Newcastle Univ.) and Ken Takai (JAMSTEC) for valuable discussion, and also thank Mr Gaku Sato and Mr Shigeru Shimamura for assisting experiments.

This work was supported by the Human Frontier Science Program (RPG0029/2020 to Y.K.), JST PRESTO (JPMJPR18K5 to Y.K.), JSPS KAKENHI (16H06156, 16KK0161, 16H00797, 26119704 to Y.K.), Astrobiology Center Project of the National Institutes of Natural Sciences (AB291017 to Y.K.), and an internal Grant-in-Aid from Earth-Life Science Institute (to Y.K.).

## Author Contributions

S.E. and R.M. performed most of the experiments, and M.F., S.B., and Y.S. assisted some basic experiments. T.K. performed the mass spectrometry analysis and interpreted the data. Y.K. designed the framework of the research, analyzed the data, and wrote the paper. All authors discussed the results and commented on the paper.

## Competing interests

The authors declare no competing interests.

## Supplementary Information

Supplementary Figures 1-17

Supplementary Supplementary Table 1-10

Supplementary Text 1-2

## Notes

### Competing Interest Statement

The authors have declared no competing interest.

